# QDeep: distance-based protein model quality estimation by residue-level ensemble error classifications using stacked deep residual neural networks

**DOI:** 10.1101/2020.01.31.928622

**Authors:** Md Hossain Shuvo, Sutanu Bhattacharya, Debswapna Bhattacharya

**Affiliations:** Department of Computer Science and Software Engineering, Auburn University, Auburn, AL 36849, USA; Department of Biological Sciences, Auburn University, Auburn, AL 36849, USA

## Abstract

**Motivation:** Protein model quality estimation, in many ways, informs protein structure prediction. Despite their tight coupling, existing model quality estimation methods do not leverage inter-residue distance information or the latest technological breakthrough in deep learning that has recently revolutionized protein structure prediction.

**Results:** We present a new distance-based single-model quality estimation method called QDeep by harnessing the power of stacked deep residual neural networks (ResNets). Our method first employs stacked deep ResNets to perform residue-level ensemble error classifications at multiple predefined error thresholds, and then combines the predictions from the individual error classifiers for estimating the quality of a protein structural model. Experimental results show that our method consistently out-performs existing state-of-the-art methods including ProQ2, ProQ3, ProQ3D, ProQ4, 3DCNN, MESHI, and VoroMQA in multiple independent test datasets across a wide-range of accuracy measures; and that predicted distance information significantly contributes to the improved performance of QDeep.

**Availability:** https://github.com/Bhattacharya-Lab/QDeep

## 1 Introduction

Estimating the quality of a computationally generated protein structural model serves as a key component of protein structure prediction (Won *et al.*, 2019; Uziela *et al.*, 2017). Model quality estimation assists in validating and evaluating predicted protein models at multiple stages of a structure prediction pipeline, thus greatly affecting its prediction accuracy (Cao *et al.*, 2015; Kalman and Ben-Tal, 2010). Methods for model quality estimation can be broadly categorized into two major classes that include “single-model” methods and “consensus” approaches. Single-model methods estimate structural quality purely from the model itself (Derevyanko *et al.*, 2018; Sato and Ishida, 2019; Uziela *et al.*, 2017, 2016, 3; Ray *et al.*, 2012, 2; Pagès *et al.*, 2019; Karasikov *et al.*, 2019; Olechnovič and Venclovas, 2017) whereas consensus approaches exploit information from other models in a pool of possible alternatives (Cheng *et al.*, 2009; McGuffin and Roche, 2010; Benkert *et al.*, 2009; Alapati and Bhattacharya, 2018). As such, performance of consensus approaches can be tremendously affected by the size and diversity of the model pool (Manavalan and Lee, 2017; Cao *et al.*, 2016; Won *et al.*, 2019), sacrificing their generality and large-scale use in standalone structure prediction systems. Single-model methods, on the other hand, are free from such limitation and can be independently employed for scoring and model selection. As a result, single-model quality estimation methods are gaining increasing attention by the community in the recent editions of Critical Assessment of techniques for protein Structure Prediction (CASP) experiments (Cheng *et al.*, 2019; Kryshtafovych *et al.*, 2018, 12; Won *et al.*, 2019), the universal standard for objectively evaluating the state of the art of protein structure prediction.

Single-model quality estimation methods use various combinations of features and employ different machine learning approaches for estimating the quality of a protein model without any knowledge of the experimental structure by learning a mapping from the features to its quality. For instance, ModelEvaluator (Wang *et al.*, 2009) uses structural features to train Support Vector Machine (SVM). RFMQA (Manavalan *et al.*, 2014) utilizes structural features and potential energy terms for training Random Forest (RF), ProQ2 (Ray *et al.*, 2012, 2) uses evolutionary sequence profile combined with contacts and other structural features to train SVM. In addition to these traditional machine learning-based methods, a growing number of approaches are employing deep learning, some of which delivering top performance in the most recent CASP13 experiment (Won *et al.*, 2019; Cheng *et al.*, 2019). For example, ProQ3D (Uziela *et al.*, 2017) and ProQ4 (Hurtado *et al.*, 2018) exploit the strengths of multi-layer perceptron and 1D fully convolutional neural network (CNN), respectively. Other recent methods such as Ornate (Pagès *et al.*, 2019), and 3DCNN (Derevyanko *et al.*, 2018; Sato and Ishida, 2019) take advantage of 3D CNNs, attaining state-of-the-art performance.

Despite their effectiveness, these approaches do not consider some key factors that can significantly improve single-model quality estimation performance. First, accurate prediction of inter-residue distance information has dramatically improved the nature of protein model generation (Senior *et al.*, 2020; Greener *et al.*, 2019; Xu, 2019), but none of the quality estimation methods incorporate distance information. Second, very deep and fully convolutional residual neural network (ResNet) (He *et al.*, 2016) has emerged as a breakthrough deep learning technology that has revolutionized many computer vision tasks and very recently protein contact or distance prediction (Li *et al.*, 2019; Wang *et al.*, 2017), but their power has not yet been harnessed in estimating model quality. Third, most of these machine leaning-based approaches typically rely on a single trained predictor for quality estimation either at the global or local level. That is, they do not make use of ensemble learning.

In this article, we present a brand-new distance-based single-model quality estimation method QDeep by training an ensemble of stacked deep ResNets. Such architecture can perform residue-level ensemble error classifications at multiple predefined error thresholds. Here we use 1, 2, 4, and 8Å as the error thresholds to model the GDT-TS score (Zemla, 2003) by predicting the likelihood of the error between the Cα atom of any residue of a model to be within rÅ from the corresponding aligned residue in the experimental structure, where r ∈ {1, 2, 4, 8}Å. Combined predictions from the individual error classifiers can then be used to estimate the quality of a protein structural model. We train QDeep using a redundancy-removed set of proteins from CASP9 and CASP10, validate the performance of the individual deep ResNet models on CASP11, and then evaluate its quality estimation performance on CASP12 and CASP13 targets. Our experimental results show that our method yields much better performance than existing methods and also results in better discrimination between “good” and “bad” models. The improved performance of QDeep is deeply driven by our effective integration of distance information for single-model quality estimation. QDeep is freely available at https://github.com/Bhattacharya-Lab/QDeep.

## 2 Materials and methods

The flowchart of QDeep is shown in Figure 1, which consists of four steps: multiple sequence alignment generation, feature collection, stacked deep ResNets training, and residue-level ensemble error classifications and their combination for model quality estimation.

**Fig. 1.**
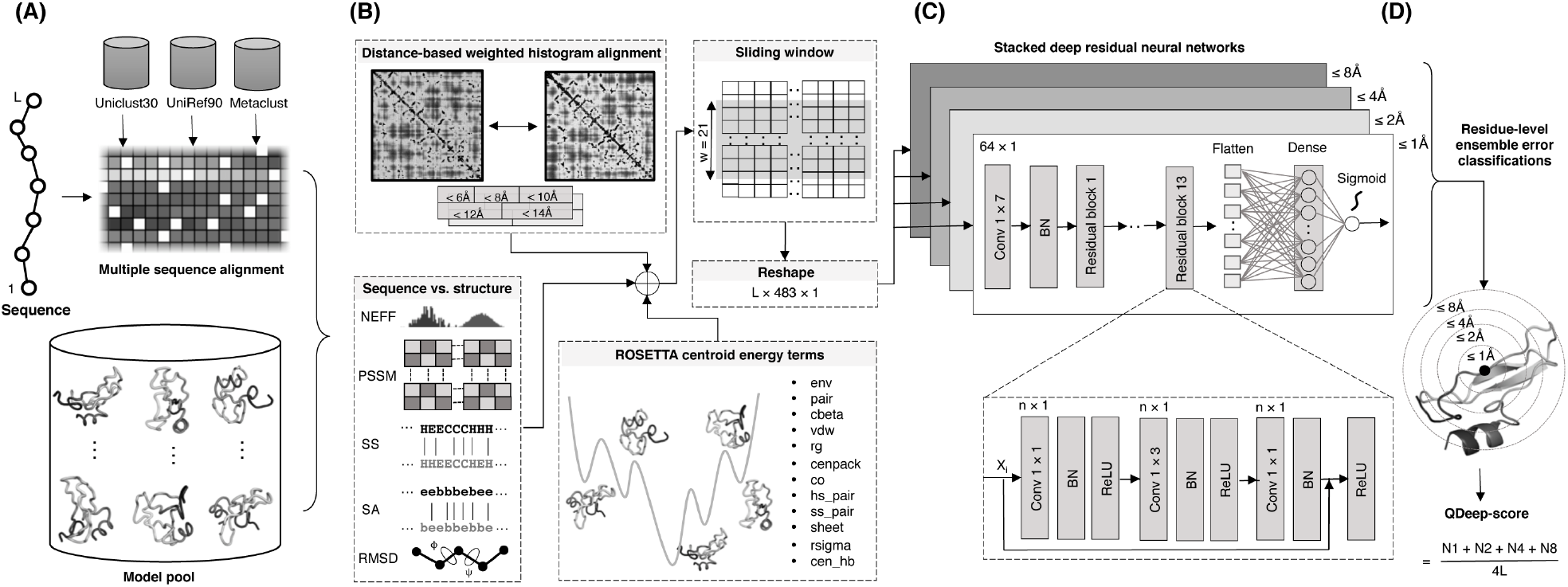
Flowchart of QDeep. (A) Multiple sequence alignment generation. (B) Distance-based, sequence vs. structure consistency-based, and ROSETTA centroid energy terms-based features collection. (C) Architecture of stacked deep residual neural network classifiers at 1, 2, 4, 8Å error thresholds (D) Residue-level ensemble error classifications and their combination for model quality estimation.

### 2.1 Multiple sequence alignment generation

We generate multiple sequence alignment (MSA) (Figure 1A) using HHblits (Remmert *et al.*, 2012) with a query sequence coverage of 10% and pairwise sequence identity of 90% against whole-genome sequence database Uniclust30 (Mirdita *et al.*, 2017) for three iterations with an Evalue inclusion threshold of 10^−3^. We also experiment the inclusion of other whole-genome sequence database UniRef90 (Suzek *et al.*, 2015) and metagenome database Metaclust (Steinegger and Söding, 2018) using the DeepMSA (Zhang *et al.*, 2019) pipeline to generate more sensitive and diverse MSA with improved coverage and alignment depth. The generated MSA serves as the key input to inter-residue distance prediction as well as other sequence-based features such as secondary structure and solvent accessibility.

### 2.2 Feature collection

As shown in Figure 1B, we generate a total of 23 features for describing each residue of a model that includes distance-based weighted histogram alignment, sequence vs. structure consistency, and ROSETTA centroid energy terms. We briefly describe them below.

#### 2.2.1 Distance-based weighted histogram alignment

We predict inter-residue distance map of the target protein by feeding the MSA to DMPfold (Greener *et al.*, 2019) and obtain the initial distance prediction without any iterative refinement (i.e., rawdistpred.current files) containing 20 distance bins with associated likelihoods between the interacting residue pairs. The distance map is then converted to 5 evenly distributed distance intervals: 6Å, 8Å, 10Å, 12Å, and 14Å by summing up likelihoods for distance bins below specific distance thresholds and considering only the residue pairs having likelihoods of at least 0.2 to reduce noise. We calculate observed inter-residue distance histogram for each model in the model pool at the same five 5 distance intervals mentioned above to perform dynamic programing alignments of the predicted and observed distance histograms through eigen-decomposition (Di Lena *et al.*, 2010). The 5 alignment scores, each of which ranges between 0 and 1, are used as 5 distance-based features after multiplying with empirically selected weights of 0.10, 0.25, 0.30, 0.25 and 0.10 for alignment scores at distance bins 6, 8, 10, 12 and 14Å, respectively, to allow higher emphasis of the distance intervals between 8 and 12Å.

#### 2.2.2 Sequence vs. structure consistency

##### Number of effective sequences

We use the normalized number of effective sequences, (NEFF) (Zhang *et al.*, 2019) as a feature. NEFF is calculated as the reciprocated sum of the number of sequences in the MSA with a sequence identity greater than 80% to the n^th^ sequence, divided by the total number of sequences in the MSA.

##### Sequence profile conservation score

We generate sequence profile by searching the NR database using PSI-BLAST v2.2.26 software (Altschul *et al.*, 1997) with an E-value of 0.001. The information per position score from the resulting position specific scoring matrix (PSSM) is then used as a feature after applying sigmoidal transformation to scale the score between 0 and 1.

##### Secondary structure and solvent accessibility

We predict secondary structure (SS) and solvent accessibility (SA) using SPIDER3 (Heffernan *et al.*, 2017) and use residue-specific binary agreement between predicted and observed secondary structure as well as the squared error between predicted and observed relative solvent accessibility as features.

##### Angular RMSD

We use normalized angular root mean square deviation (RMSD) between the two backbone dihedral angles (ϕ, Ψ) predicted from the sequence using SPIDER3 and their observed values computed from the models as two features, which are computed as:

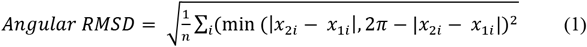

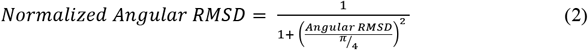

where x_1i_ is the vector of ϕ or Ψ angle sequence for n residues predicted from the amino acid sequence and x_2i_ is the vector of the corresponding observed ϕ or Ψ angle sequence for n residues in the model.

#### 2.2.3 ROSETTA centroid energy terms

We use 12 ROSETTA (Leaver-Fay *et al.*, 2011; Rohl *et al.*, 2004) centroid energy terms as features that include residue environment (env), residue pair interactions (pair), cβ density (cbeta), steric repulsion (vdw), radius of gyration (rg), packing (cenpack), contact order (co), statistical potentials for secondary structure formation (hs_pair, ss_pair, sheet, rsigma), and centroid hydrogen bonding (cen_hb). We apply sigmoidal transformations to scale the energy terms before using them as features.

#### 2.2.4 Sliding window

In order to capture the local interactions among residues, we employ a sliding window of 21 around the central residue (i.e., 10 residues on both sides) similar to (Uziela *et al.*, 2016). This results in a 483-dimensional feature vector for each residue. N- or C-terminal residues having one or more missing neighbors on either side are padded with additional 0’s to match the feature dimension.

### 2.3 Architecture of stacked deep residual neural networks

Figure 1C shows a high-level overview of the architecture of our stacked deep residual neural networks. Each network consists of 13 residual blocks with three 1-dimensional convolutional layers in each block. We adopt the bottleneck design (He *et al.*, 2016) for the residual modules with a kernel size of 1 × 1, 1 × 3, and 1 × 1, respectively, for three convolutional layers in each residual block. Here the 1 × 1 layers at the beginning and at the end of each residual block are used to reduce and restore the dimensionality of the feature vector, thus maintaining the consistency in the dimensionality of feature maps throughout the network. We input L × 483 × 1 feature to a convolutional layer with a kernel size of 1 × 7 and filter of 64 × 1. Afterwards, the feature is transformed into 64-dimensional feature vector using 1 × 1 convolutional layer of the first residual block with a filter size of 64 × 1. In each of the residual block, a “shortcut connection” between the input and the output layer, skipping the intermediate layer is established. This connection works as identity mapping and its outputs are added to the output from all previously stacked layers and passed to the pre-activation phase of the final layer of a residual block. As depicted in Figure 1C, the input x in the i^th^ layer is added to the input of the final layer of a residual block.

Therefore, the activation in the output layer for a specific residual block is applied to the x_i_ + Ƒ (x_i_+1). Formally, it is represented as:

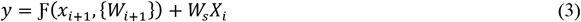

where Ƒ is the ReLU activation function, W is the weight vector in a particular layer i, W_s_ is the additional parameter to the model, representing the linear projection by the shortcut connection, applied to match the dimension which is implemented by 1 × 1 convolution.

Our entire deep residual network is divided into three stages with 3 residual blocks in the first stage, 4 in the second stage and 6 in the last stage. In Figure 1C, n defines the output channels for the residual block in each stage that are set to 128, 256 and 512, respectively. Therefore, in the first block of each stage, the feature map is halved and the filter size is increased by a factor of 2. The dimensionality of the feature map remains the same in the consecutive blocks in a stage. We apply batch normalization on the input features before passing to the convolutional layer in the first stage of the residual network. The utilization of this setting, therefore, minimizes the internal covariate shift as well as the need for the Dropout (Ioffe and Szegedy, 2015).

At the end of the residual blocks, we use an average pooling layer with a pool size of 2 that reduces the number of parameters and helps in faster computation. Finally, a flatten layer accepts the pooled feature map to transform into a 1D vector and passes to the fully connected (i.e., dense) layer.

### 2.4 Model training

We collect submitted models from CASP9 and CASP10 (Moult *et al.*, 2016, 2018) experiments for a total of 220 protein targets, whose experimental structures are publicly available. On an average, there are 282 models per target. To remove redundancy, we perform MUFOLD clustering (Zhang and Xu, 2013) and select the centroid of the top 10 clusters. It should be noted that not all the targets have all 10 clusters and MUFOLD fails to execute for 5 targets, resulting in a total of 2,130 redundancy-removed models having 303,675 residues for 215 CASP targets. We prepare four sets of features at 1, 2, 4, and 8Å error thresholds with the same training data by assigning a binary label to each one of the residues after calculating the errors between the Cα atoms of each of the residues in the model and the corresponding aligned residue in the experimental structure using the LGA program (Zemla, 2003). We assign a label of 1 (positive class) to the feature set of each of the residues, if the error is within rÅ (where r ∈ {1, 2, 4, 8}Å following GDT-TS), 0 (negative class) otherwise.

We train an ensemble of four independent ResNet models using the four sets of features at 1, 2, 4, and 8Å error thresholds. As Conv1D accepts an input shape of a 3D tensor with batch, steps, and channel, respectively, we reshape the feature vector into L × 483 × 1 prior to passing to the input layer. We train the networks with the maximum number of epochs of 120 and an optimal batch size of 64 that best fits the GPU limit. Also, to avoid overfitting we use EarlyStopping callback of Keras (Chollet, 2015) with a patience value of 20. We optimize the model using the binary crossentropy loss function and first-order gradient-based Adam optimizer (Kingma and Ba, 2014).

### 2.5 Residue-level ensemble error classifications and their combination for model quality estimation

Figure 1D shows the residue-level ensemble classifications and their combination for model quality estimation. We consider each one of the four deep ResNet models as an independent residue-specific error classifier, while their ensemble collectively estimates the structural quality of a model. For each classifier, the output layer with the sigmoid activation function predicts the likelihood of residue-level errors at 1, 2, 4, and 8Å error thresholds. We set a likelihood cutoff of 0.5 to convert the likelihood of a residue-level error to binary classification, where a likelihood value greater than the cutoff is classified as 1, indicating the residue-specific error to be within rÅ error level (r ∈ {1, 2, 4, 8}Å) and 0 otherwise. For a given error threshold at rÅ, we calculate the aggregated error for the whole model by summing up the number of residues belonging to the positive class N_r_. Analogous to GDT-TS score, we estimate the quality of a model by combining the ensemble of residue-level classifiers as:

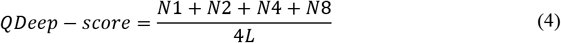

where N1, N2, N4, and N8 are the number of aligned residues within 1, 2, 4 and 8Å error thresholds, respectively, and L is the length of the target protein. Consequently, QDeep-score lies between [0, 1] with a higher value indicating better quality.

### 2.6 Evaluation method and programs to compare

We validate the individual residue-level classifiers using the “stage 2” model pool (150 models/target) for 82 CASP11 targets (Kryshtafovych *et al.*, 2016) with publicly available experimental structures. For evaluating model quality estimation performance, we use the stage 2 model pool for 40 and 20 targets from CASP12 and CASP13, respectively, (Kryshtafovych *et al.*, 2016; Cheng *et al.*, 2019) with a total of 9,000 models for both datasets. The training and test datasets are nonoverlapping with an average pairwise sequence identity of 21%. We use three evaluation criteria to measure the performance of model quality estimation: (i) ability to reproduce the true model-native similarity scores, (ii) ability to find the best model, and (iii) ability to distinguish between good and bad models. For the first criterion, we use average per-target and global Pearson, Spearman and Kendall’s Tau correlations between the estimated scores and the true GDT-TS accuracy considering all models in a given dataset. Consequently, a higher correlation indicates better performance. For the second criterion, we use average GDTTS loss that is the difference between the true GDT-TS of the top model selected by the estimated score and that of the most accurate model in the pool. A lower loss, therefore, indicates better performance. For the third criterion, we preform receiver operating characteristics (ROC) analysis using a cutoff of GDT-TS = 0.4 to separate good and bad models. Meanwhile, the area under ROC curves (AUC) quantifies the ability of a method to distinguish good and bad models.

We compare our new distance-based single-model method QDeep with state-of-the-art single-model quality estimation methods that include ProQ2 (Ray *et al.*, 2012, 2), ProQ3 (Uziela *et al.*, 2016, 3), ProQ3D (Uziela *et al.*, 2017), ProQ4 (Cheng *et al.*, 2019, 13; Hurtado *et al.*, 2018), 3DCNN (Sato and Ishida, 2019), MESHI (Kalisman *et al.*, 2005), and VoroMQA (Olechnovič and Venclovas, 2017). For CASP12 targets, all tested methods are run locally with parameters set according to their respective papers. For CASP13, we directly obtain quality estimation predictions submitted by the tested methods from the data archive of the CASP official website.

## 3 Results

### 3.1 Validation of individual residue-level classifiers

We validate the performance of our individual deep ResNet-based classifiers on 82 CASP11 targets. Figure 2 shows the ROC curves for each of the classifiers trained at error thresholds of 1, 2, 4, and 8Å, respectively. All individual classifiers achieve AUC values ~0.8, demonstrating their effectiveness at various error thresholds. Of note, the AUC values steadily increase at higher values of error thresholds. This is not surprising because at a lower error threshold, the proportion of residues belonging to the positive class is very low. That is, the number of positive and negative labels is extremely unbalanced. For example, in the training dataset the ratios between the positive and negative labels are 0.27 (88,936/331,318), 0.54 (147,194/273,084), 0.93 (202,373/217,927) and 1.57 (256,740/163,566) for 1, 2, 4, and 8Å error thresholds, respectively. The sparsity of positive labels at lower error thresholds may be the reason behind their somewhat lower performance. Nonetheless, they still deliver reasonable residue-level classification performance. Binary classification performance metrics such as F1, MCC, Precision, and Recall for the individual classifiers at various error thresholds are reported in Supplementary Table S1.

**Fig. 2.**
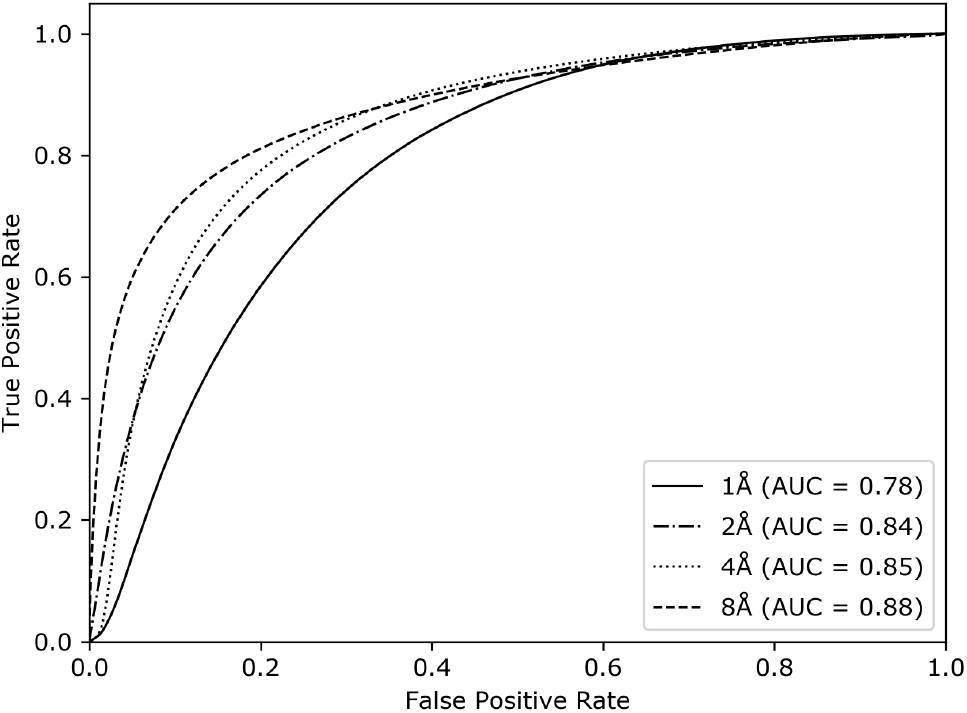
Accuracy of the individual residue-level classifiers at 1, 2, 4 and 8Å error thresholds on the validation set of 82 CASP11 targets.

### 3.2 Performance evaluation on CASP datasets

Table 1 reports the performance of our new method QDeep and five other top-performing single-model quality estimation methods on CASP12 and CASP13 stage 2 datasets. QDeep consistently outperforms all other tested methods for both CASP12 and CASP13 sets across almost all performance criteria. For instance, QDeep attains the highest per-target average Pearson correlation of 0.740 in CASP12, which is much better than the second-best ProQ3D (0.688). Additionally, QDeep attains the lowest average GDT-TS loss of 0.051, which is significantly lower than the second-best ProQ3 (0.071). Furthermore, QDeep always delivers the highest global correlations in CASP12. The same trend continues for CASP13 set, in which QDeep attains the highest per-target average Pearson correlation of 0.752, better than the second-best ProQ4 (0.733). In terms of GDT-TS loss in CASP13, however, MESHI attains the lowest average GDT-TS loss (0.070) as it adopts an additional loss-enrichment step in its pipeline. QDeep has an unusually high GDT-TS loss of 0.455 for the CASP13 target T1008, which raises its average GDT-TS loss. On further inspection, we find that the alignment depth of the MSA for T1008 is zero having no identifiable homologous sequences. If T1008 is excluded, the average loss of QDeep becomes 0.068. Considering all targets though, the average GDT-TS loss of QDeep in CASP13 is still comparable to the other tested methods. The global correlations attained by QDeep are always the highest in CASP13. Of note, ProQ4, the second-best performing method in CASP13 after QDeep in terms of per-target average correlations, consistently exhibits poor global correlations. MESHI, the method attaining the best average GDT-TS loss in CASP13, does not deliver top performance in terms of per-target average correlations. That is, there are complementary aspects of model quality estimation that can lead to performance trade-offs. Our new method QDeep strikes an ideal balance to deliver top-notch performance across various facets of model quality estimation simultaneously.

**Table 1.**
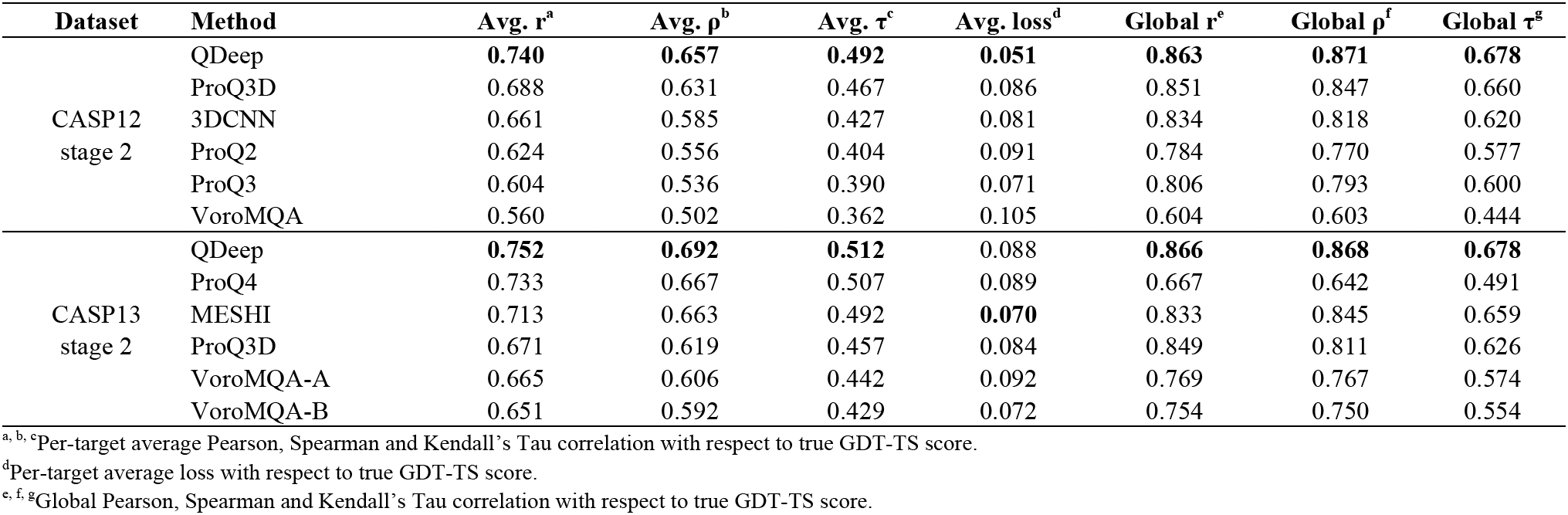
Performance of single-model quality estimation methods on CASP12 and CASP13 stage 2 datasets, sorted in decreasing order of average pertarget Pearson correlations. Values in bold represent the best performance.

| Dataset | Method | Avg. $r^a$ | Avg. $\rho^b$ | Avg. $\tau^c$ | Avg. loss<sup>d</sup> | Global $r^e$ | Global $\rho^f$ | Global $\tau^g$ |
| --- | --- | --- | --- | --- | --- | --- | --- | --- |
| CASP12 stage 2 | QDeep | **0.740** | **0.657** | **0.492** | **0.051** | **0.863** | **0.871** | **0.678** |
|  | ProQ3D | 0.688 | 0.631 | 0.467 | 0.086 | 0.851 | 0.847 | 0.660 |
|  | 3DCNN | 0.661 | 0.585 | 0.427 | 0.081 | 0.834 | 0.818 | 0.620 |
|  | ProQ2 | 0.624 | 0.556 | 0.404 | 0.091 | 0.784 | 0.770 | 0.577 |
|  | ProQ3 | 0.604 | 0.536 | 0.390 | 0.071 | 0.806 | 0.793 | 0.600 |
|  | VoroMQA | 0.560 | 0.502 | 0.362 | 0.105 | 0.604 | 0.603 | 0.444 |
| CASP13 stage 2 | QDeep | **0.752** | **0.692** | **0.512** | 0.088 | **0.866** | **0.868** | **0.678** |
|  | ProQ4 | 0.733 | 0.667 | 0.507 | 0.089 | 0.667 | 0.642 | 0.491 |
|  | MESHI | 0.713 | 0.663 | 0.492 | **0.070** | 0.833 | 0.845 | 0.659 |
|  | ProQ3D | 0.671 | 0.619 | 0.457 | 0.084 | 0.849 | 0.811 | 0.626 |
|  | VoroMQA-A | 0.665 | 0.606 | 0.442 | 0.092 | 0.769 | 0.767 | 0.574 |
|  | VoroMQA-B | 0.651 | 0.592 | 0.429 | 0.072 | 0.754 | 0.750 | 0.554 |

<sup>a, b, c</sup>Per-target average Pearson, Spearman and Kendall's Tau correlation with respect to true GDT-TS score.
<sup>d</sup>Per-target average loss with respect to true GDT-TS score.
<sup>e, f, g</sup>Global Pearson, Spearman and Kendall's Tau correlation with respect to true GDT-TS score.

To investigate the ability of QDeep to distinguish good and bad models in comparison with the other tested methods, we perform ROC analysis using all models for all targets in CASP12 and CASP13. Figures 3 shows the ROC curves with AUC values. Once again, QDeep consistently achieves the highest AUC values for both CASP12 and CASP13. The AUC of QDeep is slightly higher than the second-best ProQ3D in CASP12 and noticeably higher in CASP13, demonstrating its better performance in separating good and bad models compared to the others.

**Fig. 3.**
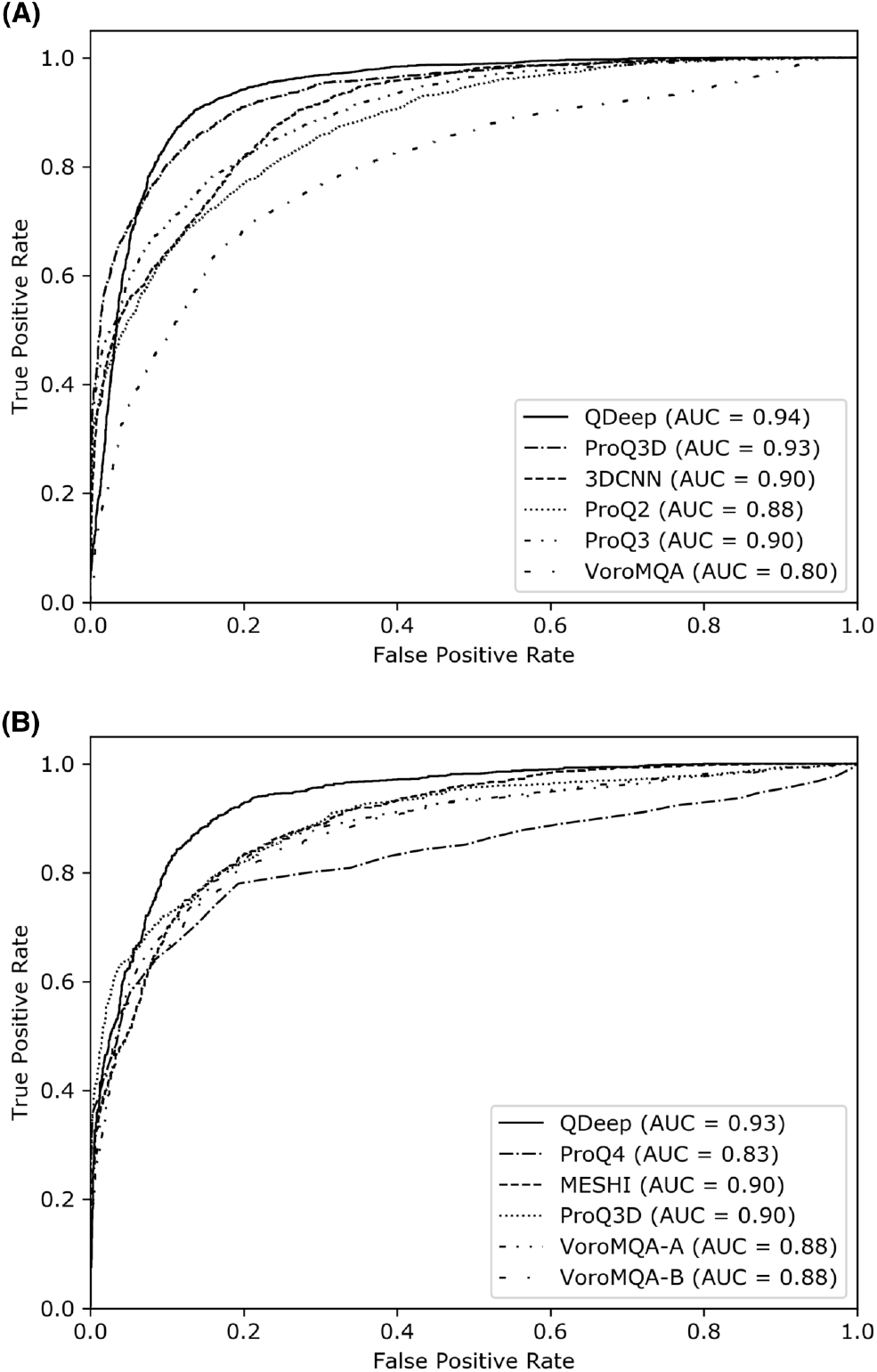
The ability of single-model quality estimation methods to distinguish good and bad models in (A) CASP12 and (B) CASP13 stage 2 datasets. A cutoff of GDT-TS = 40 is used to separate good and bad models.

It is interesting to note that among the other tested methods, deep learning-based approaches such as ProQ4, ProQ3D, and 3DCNN routinely deliver better performance in both CASP12 and CASP13 datasets. Among the ProQ series of predictors, deep learning-based methods such as ProQ3D and ProQ4 perform better than SVM-based approaches like ProQ2 and ProQ3. While these methods themselves are intrinsically different making it difficult if not impossible to firmly conclude the underlying cause of their performance differences, the trend of superior performance of deep learning-based methods may indicate the inherent advantage of transitioning from traditional machine learning to deep learning-based approaches for model quality estimation task. To investigate the effect of various deep architectures on quality estimation performance when everything else remains the same, we compare deep ResNet with Long Short-Term Memory (LSTM) (Hochreiter, Sepp, 1997) and Convolutional Neural Network (CNN) (Lee *et al.*, 2009) by performing controlled experiments. Equivalent to the ResNet model ensemble employed in QDeep, we train ensembles of four independent LSTM and CNN residue-specific error classifiers at 1, 2, 4, and 8Å error thresholds using the same features sets and same training data used in QDeep. The model architectures and training procedure for the LSTMs and CNNs are described in Supplementary Method. Table 2 presents the head-to-head performance comparison between ResNets, LSTMs, and CNNs on CASP12 and CASP13 stage 2 datasets. The results show that ResNet delivers the best performance across all performance criteria, while LSTM consistently outperforms CNN. ResNet attains the highest per-target average correlations and the lowest average GDT-TS losses in both CASP12 and CASP13 sets. LSTM attains an average GDT-TS loss of 0.059, which is lower than all tested methods in CASP12 and second only to the ResNet architecture of QDeep. In CASP13, LSTM delivers better per-target average Pearson (0.735) and Spearman (0.668) correlations than all other methods except ResNet-based QDeep. The results further emphasize the advantage of using sophisticated deep architecture such as LSTM for model quality estimation. Our new method QDeep goes one step further by training an ensemble of state-of-the-art stacked deep ResNets classifiers, delivering the best predictive performance. Meanwhile, the novel use of distance information in QDeep substantially improves the performance, as discussed later. In summary, the advantage of QDeep in single-model quality estimation over the others is manifold.

**Table 2.** Performance comparison of deep ResNet models used in QDeep with other deep learning architectures on CASP12 and CASP13 stage 2 datasets. Values in bold represent the best performance.

|  | CASP12 stage 2 |  |  |  | CASP13 stage 2 |  |  |  |
| --- | --- | --- | --- | --- | --- | --- | --- | --- |
| | Avg. $r^a$ | Avg. $\rho^b$ | Avg. $\tau^c$ | Avg. loss<sup>d</sup> | Avg. $r^a$ | Avg. $\rho^b$ | Avg. $\tau^c$ | Avg. loss<sup>d</sup> |
| ResNet | **0.740** | **0.657** | **0.492** | **0.051** | **0.752** | **0.692** | **0.512** | **0.088** |
| LSTM | 0.716 | 0.596 | 0.452 | 0.059 | 0.735 | 0.668 | 0.500 | 0.116 |
| CNN | 0.657 | 0.581 | 0.433 | 0.097 | 0.735 | 0.660 | 0.487 | 0.116 |

The legends for columns 2-9 convey the same meaning as mentioned in Table 1.

### 3.3 Impact of deeper sequence alignment

Since a large number of features used in QDeep is dependent on MSA, which has been shown to significantly affect contact prediction, secondary structure prediction, and threading (Zhang *et al.*, 2019), we replace our alignment generation component with DeepMSA (Zhang *et al.*, 2019) to generate deeper sequence alignments by integrating wholegenome and metagenome sequence databases and retrain the ensemble of four stacked deep ResNet models with the same architecture and training procedure mentioned earlier (hereafter called QDeep^DeepMSA^). To study the impact of deeper MSA in model quality estimation, we perform head-to-head performance comparison between QDeep and QDeep^DeepMSA^. To make a fair comparison, we use the same test datasets of CASP12 and CASP13 with the same feature sets. As reported in Table 3, QDeep^DeepMSA^ further improves per-target average correlations in both datasets. The improvement is particularly noticeable for CASP13, in which QDeep^DeepMSA^ attains average per-target Pearson, Spearman, and Kendall’s Tau correlations of 0.777, 0.720, and 0.538, respectively, that are much higher than QDeep trained on standard MSA. In terms of average GDT-TS loss, QDeep^DeepMSA^ results in slight improvement in CASP13 but visible worsening in CASP12. In Supplementary Figure S1, we show the performance of the individual classifiers at 1, 2, 4, and 8Å error thresholds by performing ROC analysis on our validation set comprising of 82 CASP11 targets. Except 8Å error threshold, all other deep ResNet classifiers trained on deeper alignments attain higher AUC values compared to their counterparts using standard MSA. That is, deeper sequence alignments can be advantageous to further improve the performance of our new quality estimation method QDeep, particularly by enhancing its ability to better reproduce true model-native similarity scores. To understand whether it is possible to further improve the performance by finetuning the hyperparameters of the ResNet architecture to better leverage the deeper sequence alignments, we train shallower and deeper ResNet architectures. For the shallower architecture, we stack 2 blocks in each of the three stages, resulting in a total of 6 residual blocks for each independent residue-level error classifier. The deeper architecture consists of a total of 20 residual blocks for each independent residue-level error classifier having 6, 7, and 7 blocks sequentially for the three stages. As described in Supplementary Methods, these two variants of our original 13-residual-blocks-QDeep^DeepMSA^ are trained using the same features with deep MSAs and same training data. Head-to-head performance comparison reported in Supplementary Table S2 reveals minor performance variations between the variants, with all architectures outperforming the other tested methods for both CASP12 and CASP13 sets in majority of performance criteria. The deeper architecture with 20 residual blocks performs better than the shallower architecture with 6 residual blocks, particularly for CASP13. Our original ResNet architecture of QDeep^Deep-^ MSA consistently delivers the best performance over the variants across all assessment metrics, validating its effectiveness for quality estimation.

**Table 3.** Performance comparison of variants of QDeep on CASP12 and CASP13 stage 2 datasets.

|  | CASP12 stage 2 |  |  |  | CASP13 stage 2 |  |  |  |
| --- | --- | --- | --- | --- | --- | --- | --- | --- |
| | Avg. $r^a$ | Avg. $\rho^b$ | Avg. $\tau^c$ | Avg. loss<sup>d</sup> | Avg. $r^a$ | Avg. $\rho^b$ | Avg. $\tau^c$ | Avg. loss<sup>d</sup> |
| QDeep | 0.740 | 0.657 | 0.492 | 0.051 | 0.752 | 0.692 | 0.512 | 0.088 |
| QDeep<sup>DeepMSA</sup> | 0.741 | 0.667 | 0.505 | 0.062 | 0.777 | 0.720 | 0.538 | 0.084 |
| QDeep<sup>NoDistance</sup> | 0.677 | 0.601 | 0.442 | 0.065 | 0.668 | 0.613 | 0.445 | 0.091 |

The legends for columns 2-9 convey the same meaning as mentioned in Table 1.
QDeep<sup>DeepMSA</sup>: QDeep classifiers retrained using features generated by integrating deep MSA.
QDeep<sup>NoDistance</sup>: QDeep classifiers retrained without using any distance-based features, but utilizing deep MSA.

### 3.4 Contribution of distance information

To evaluate the contribution of distance information in QDeep, we retrain an extra set of the same ensemble classifiers at 1, 2, 4, and 8Å error thresholds after excluding distance-based features, while still utilizing deep MSA for feature generation (hereafter called QDeep^NoDistance^). Head-to-head performance comparisons on the same test sets of CASP12 and CASP13 datasets reveal that QDeep^NoDistance^ performs much worse than the original distance-based QDeep method, let alone its variant QDeepDeepMSA trained using deep alignments and distance information. As shown in Table 3, QDeep^NoDistance^ substantially degrades per-target average Pearson, Spearman, and Kendall’s Tau correlations in both CASP12 and CASP13 sets. There is also a noticeable increase in the average GDT-TS loss. Clearly, the exclusion of distance information negatively affects the estimation of model quality performance. The results underscore the importance of incorporating distance information in single-model quality estimation methods such as QDeep.

## 4 Conclusion

This paper presents QDeep, a new distance-based single-model protein quality estimation method based on residue-level ensemble error classifications using stacked deep ResNets. Experimental results show that QDeep works much better than existing approaches across various accuracy measures of model quality estimation. QDeep outperforms not only currently popular ProQ series of methods including its most recent editions ProQ3D and ProQ4 but also top-ranked single-model quality estimation methods participating in the most recent 13th edition of CASP. Among the competing methods, deep learning-based approaches show a general trend of superior performance. Our new method QDeep takes a leap forward by employing cutting-edge deep learning architecture to effectively integrate predicted distance information with other sequential and structural features, leading to improved performance.

Different from the other state-of-the-art methods such as ProQ4 and MESHI that focus only on some aspects of quality estimation, our method works well on a wide-range of accuracy metrics to deliver an overall well-rounded performance. Controlled experiment on multiple datasets confirms that the improved performance of QDeep is primarily attributed to our effective integration of distance information that can be further improved in part by incorporating deeper sequence alignments. This should make distance-based protein model quality estimation a promising new avenue for many more single-model methods in the near future.

## Acknowledgements

This work was made possible in part by a grant of high performance computing resources and technical support from the Alabama Supercomputer Authority.

## Funding

This work has been partially supported by an Auburn University new faculty start-up grant to DB.

*Conflict of Interest:* none declared.

## Notes

### Competing Interest Statement

The authors have declared no competing interest.

