## Supplementary Information for "QDeep: distance-based protein model quality estimation by residue-level ensemble error classifications using stacked deep residual neural networks"

### Supplementary Information
*for*

#### QDeep: distance-based protein model quality estimation by residue-level ensemble error classifications using stacked deep residual neural networks

Md Hossain Shuvo[1], Sutanu Bhattacharya[1], Debswapna Bhattacharya[1,2,*]

[1]Department of Computer Science and Software Engineering and [2]Department of Biological Sciences, Auburn University, Auburn, AL 36849, USA.

**Supplementary Table S1.** Performance of individual classifiers at 1, 2, 4, and 8Å error thresholds on 82 CASP11 targets.

| Dataset | r[a] | F1[b] | MCC[c] | Precision | Recall |
|---|---|---|---|---|---|
| CASP11 | 1Å | 0.49 | 0.33 | 0.51 | 0.48 |
| | 2Å | 0.70 | 0.52 | 0.74 | 0.66 |
| | 4Å | 0.81 | 0.57 | 0.76 | 0.85 |
| | 8Å | 0.85 | 0.56 | 0.83 | 0.88 |

[a]Deep ResNet classifier trained at a distance threshold of r
[b]F1 score
[c]Matthew's correlation coefficient

**Supplementary Figure S1.** Accuracy of the individual residue-level classifier at 1, 2, 4 and 8Å error thresholds on 82 CASP11 targets. The classifiers are trained using the features generated by integrating deep MSA.

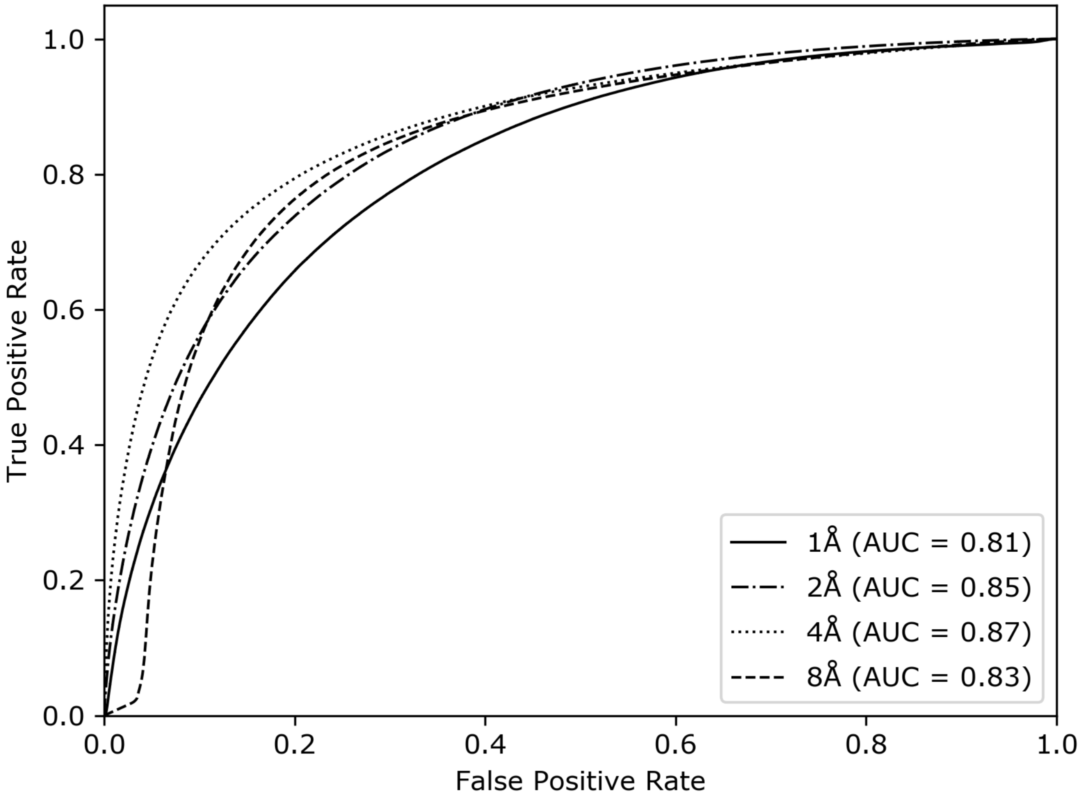

**Supplementary Methods**

#### 1. Architectures of various deep learning models for model quality estimation

To investigate the effect of various deep learning models on quality estimation performance when everything else remains the same, we train Long Short-Term Memory (Hochreiter, Sepp, 1997) and Convolutional Neural Networks (Lee *et al.*, 2009) in addition to the deep ResNet architecture used in QDeep.

##### 1.1. Long Short-Term Memory (LSTM):

We construct an LSTM network by stacking LSTM layer with 64 LSTM blocks and two subsequent hidden dense layers with 32 and 16 nodes, respectively. Similar to the deep ResNet model ensemble employed in QDeep, we train ensembles of four independent LSTM residue-specific error classifiers at 1, 2, 4, and 8Å error thresholds using the same features sets and same training data used in QDeep. We use "*relu*" activation function for each of the dense layers, and "*sigmoid*" activation function for the output layer to return a probability value between 0 and 1 for each of the classes by using a probability threshold of 0.5 for each of the predictors to classify the residue-level error to be within rÅ (where $r \in \{1, 2, 4, 8\}$Å).

##### 1.2. Convolutional Neural Network (CNN):

We construct a CNN architecture having 5 sequential stacked layers. We use 64 filters for each of the layers with a kernel size of 3 and "relu" activation function. To reduce the feature dimensionality, we add a *MaxPooling1D* layer with a pool size of 2 after every two convolutional layers. Additionally, to scale the activation we use BatchNormalization layer after every 2 convolutional layers. To reduce overfitting, we add a *Dropout* layer with a 50% dropout after every 2 convolutional layers. Subsequently, we add a *Flatten* layer followed by a fully connected dense layer as an output layer. The *Flatten* layer transforms the feature into 1D vector before it is passed to the output layer. Once again, we train ensembles of four independent CNN residue-specific error classifiers at 1, 2, 4, and 8Å error thresholds using the same features sets and same training data used in QDeep. We use "*sigmoid*" activation function in the dense layer to return a probability value between 0 and 1 for each of the classes by using a probability threshold of 0.5 for each of the predictors to classify the residue-level error to be within rÅ (where $r \in \{1, 2, 4, 8\}$Å).

#### 2. Training various architectures of deep residual neural networks (ResNets)

In addition to the original deep ResNet model employed in the QDeep method that consists of 13 residual blocks (hereafter called $\text{ResNet}^{B13}$), we investigate the effect of the hyperparameters of the ResNet architecture on quality estimation performance by training additional ResNet models. We vary the number of residual blocks to train two variants of the original ResNet architecture: one shallower ResNet having 6 residual blocks for each independent residue-level error classifier (hereafter called $\text{ResNet}^{B6}$) and the other deeper ResNet having 20 residual blocks for each independent residue-level error classifier (hereafter called $\text{ResNet}^{B20}$). For $\text{ResNet}^{B6}$, we stack 2 blocks in each of the three stages, whereas for $\text{ResNet}^{B20}$, we stack 6, 7 and 7 blocks sequentially for the three stages. Similar to the original ResNet$^{B13}$ architecture, we adopt the same bottleneck design (He *et al.*, 2016) for ResNet$^{B6}$ and ResNet$^{B20}$ architectures. We use the same set of features powered by deep multiple sequence alignments and the same training data for training. For the additional ResNet architectures of ResNet$^{B6}$ and ResNet$^{B20}$, we independently train ensemble of four stacked deep ResNet models for residue-level error classifications at 1, 2, 4, and 8Å error thresholds using the same training method as used for our original ResNet$^{B13}$ (QDeep) architecture.

**Supplementary Table S2.** Performance comparison of various ResNet architectures on CASP12 and CASP13 stage 2 datasets. Values in bold represent the best performance.

| | **CASP12 stage 2** | | | | **CASP13 stage 2** | | | |
|---|---|---|---|---|---|---|---|---|
| | Avg. r[a] | Avg. ρ[b] | Avg.τ[c] | Avg. loss[d] | Avg. r[a] | Avg. ρ[b] | Avg. τ[c] | Avg. loss[d] |
| ResNet$^{B6}$ | 0.722 | 0.640 | 0.482 | 0.070 | 0.710 | 0.656 | 0.484 | 0.109 |
| ResNet$^{B13}$(QDeep) | **0.741** | **0.667** | **0.505** | **0.062** | **0.777** | **0.720** | **0.538** | **0.084** |
| ResNet$^{B20}$ | 0.716 | 0.637 | 0.478 | 0.065 | 0.742 | 0.684 | 0.507 | 0.103 |

[a, b, c]Per-target average Pearson, Spearman and Kendall's Tau correlation with respect to true GDT-TS score.
[d]Per-target average loss with respect to true GDT-TS score.
